# Sex-specific association between emotion negativity and neural processing of reward feedback in young children

**DOI:** 10.1101/2022.06.20.496831

**Authors:** Yu Sun Chung, Armen Bagdasarov, Michael S. Gaffrey

## Abstract

Understanding whether sex-related differences in the associations between emotion negativity and reward processing are present in early childhood has the potential to critically inform the etiology and prevention of psychiatric disorders characterized by dysregulated negative affect. Previous research suggests that altered neural processing of reward is associated with elevated levels of negative affect in adolescents and that the patterns of these associations may show sex-related differences. However, few studies have investigated the relationship between negative affect and reward processing in young children and none have directly examined potential sex-related differences in this relationship during early childhood. The current study investigated potential sex-related differences in neural response to reward feedback using EEG and their sex-specific association with emotional negativity in 140 children 4-7 years of age. Children completed a developmentally appropriate reward processing task while EEG data was recorded. Temporospatial Principal Component Analysis was used to separate overlapping feedback-related ERP components, revealing that males had greater loss-related feedback negativity and gain- and loss-related P200 values than females. At the individual level, males also showed a negative correlation between emotional negativity scores and gain-related P200 amplitudes. In line with prior work in adults for sex differences in attentional biases to emotional stimuli, these results suggest that neural mechanisms leading to sex differences in abnormal development of emotional negativity may emerge during early childhood and grow in magnitude during adolescence.

## 1. Introduction

Emotional negativity has been associated with the concurrent diagnosis or later development of a number of psychiatric disorders with known sex differences, including attention-deficit/hyperactivity disorder, oppositional defiant disorder, anxiety, and depression (Brotman et al., 2006; Copeland, Angold, Costello, & Egger, 2013; Copeland, Shanahan, Egger, Angold, & Costello, 2014; Dougherty et al., 2014; Ezpeleta, Granero, de la Osa, Trepat, & Domènech, 2016; Pagliaccio, Pine, Barch, Luby, & Leibenluft, 2018; Stringaris, Cohen, Pine, & Leibenluft, 2009; Wakschlag et al., 2015; Wakschlag et al., 2018). However, few studies have investigated the association between emotional negativity and early neural processing of reward and none to our knowledge have directly examined potential sex-related differences in this relationship during early childhood. This is a critical knowledge gap as dysregulated emotional negativity during early childhood has emerged as a highly promising transdiagnostic psychiatric symptom (Brotman et al., 2006; Stringaris, Zavos, Leibenluft, Maughan, & Eley, 2012; Wakschlag et al., 2012) that is predictive of later psychopathology (Beauchaine, 2001; Copeland et al., 2014; Deveney et al., 2015; Vogel, Jackson, Barch, Tillman, & Luby, 2019). Given that the association between early emotional negativity and later psychopathology has been found to be moderated by sex (Dougherty et al., 2015; Vogel et al., 2019), it is critical to understand whether associations between emotional negativity and neural markers of reward processing are sex-dependent in young children.

There is increasing interest in the mechanisms of emotional negativity in children as irritability is listed as a descriptor of a variety of different categories in the Diagnostic and Statistical Manual of Mental Disorders (e.g., “irritable mood” in depression; angry/irritable mood in disruptive mood dysregulation disorder, irritability in generalized anxiety disorder (Rao, 2014; Safer, 2009; Terlizzi & Villarroel, 2020)). Irritability is both common and impairing in many child and adult psychiatric disorders including externalizing and internalizing disorders (Safer, 2009; Snaith & Taylor, 1985), many of which have been found to have sex differences in disease risk, prevalence and expression of clinical symptoms (Loso et al., 2021; Seney & Sibille, 2014). Even though sex differences in the incidence of mood disorders are higher during adolescence, increasing evidence suggests that the sex difference in depression emerges during early childhood and grows in magnitude during adolescence (Breslau et al., 2017). For example, in a study of preadolescent children (ages 9-10 years, n = 11,384), males had higher scores and a greater frequency of clinically meaningful levels of psychopathology including depression and anxiety (Loso et al., 2021). According to a population-based study using large longitudinal community samples, young adults with a history of childhood disruptive mood dysregulation disorder had elevated rates of anxiety and depression compared to those with no history of psychiatric disorders during childhood or adolescence (Copeland et al., 2014). Despite the ubiquity of emotional negativity across childhood psychiatric disorders characterized by altered reward processing, including depression and anxiety (Prenoveau et al., 2010; Watson & Naragon-Gainey, 2010), the fundamental neural mechanism linking the relationship between emotional negativity and reward processing in children has been largely unknown. Some evidence indicates that children with emotional negativity show altered selective attention during emotional attention tasks (Deveney et al., 2013; Deveney et al., 2020; Rich et al., 2007). In another study with adults, higher irritability was associated with smaller reward-related feedback processing (Deveney, 2019). In an electroencephalogram (EEG) study (Deveney et al., 2020), there were irritability-related attentional biases toward negative emotion stimuli in young children (ages 4-7 years) during an emotional dot-probe, attention task. As a result, this prior work raises the possibility that negative emotionality may be related to multiple neural markers of reward processing, including attention orienting, reactivity, and subsequent processing.

Neurophysiological mechanisms supporting the evaluation of reward-related feedback processing have a long history of being studied using event-related potentials (ERPs) given their high degree of temporal resolution, which allows for a tight coupling between outcome onset and subsequent neurophysiological response. The most extensively examined ERP component associated with reward processing is the feedback-related negativity (FN) or reward positivity (RewP; i.e., gain-minus reward contrast) ERP, which is a negative or positive deflection occurring approximately 250-350 ms post-feedback at frontocentral EEG electrode sites. The accumulated evidence from both functional Magnetic Resonance Imaging (fMRI) and EEG studies indicates associations between blunted FN responding to reward feedback and severe depression symptoms in adolescents (O’Callaghan & Stringaris, 2019) and adults (Keren et al., 2018). However, few studies examined the potential link of emotional negativity, a promising transdiagnostic factor of depression in children, reward processing, and sex differences in this relationship.

The FN/RewP ERP component is usually preceded by a positive ERP component observed at fronto-central sites and peaking approximately 200ms after feedback onset: the P200 (Sallet, Camille, & Procyk, 2013). The P200 has been associated with early attention allocation to emotional stimuli (Carretié, Mercado, Tapia, & Hinojosa, 2001). The P200 is particularly involved in negative attentional bias, as amplitudes at this peak are higher in response to negative stimuli than positive ones (Carretié et al., 2001). More recently, empirical data shows that the observed P200 could reflect an early effect of the salience (rather than the valence) of stimuli (San Martín, Manes, Hurtado, Isla, & Ibañez, 2010). In fact, the P200 has been reported to be consistently associated with greater arousal levels (Carretié et al., 2001; Schutter, de Haan, & van Honk, 2004) and attention capturing by target stimuli (Potts, 2004; Potts, Martin, Burton, & Montague, 2006), supporting the idea that neural processes supporting attention facilitation to salient stimuli may be represented by the P200 (see (Kotz & Paulmann, 2011) for a review). Interestingly, along with several behavioral and ERP studies reporting sex differences in attention allocation during processing of emotional stimuli in adults (Kraines, Kelberer, & Wells, 2017; Pfabigan, Lamplmayr-Kragl, Pintzinger, Sailer, & Tran, 2014; Stoet, 2017; van Hooff, Crawford, & van Vugt, 2011), emerging evidence indicates sex differences in the P200 ERP components during several cognitive processes (e.g., (Aerts, van Mierlo, Hartsuiker, Santens, & De Letter, 2015), with males having a larger fronto-central P200 peak amplitude and a shorter P200 latency period compared to females (Bourisly & Shuaib, 2018).

The P300, corresponding to a large positive deflection that is maximal around 300-400 ms at central and posterior parietal scalp electrodes following feedback onset, is another feedback-related ERP response that overlaps in time with the FN/RewP and is also sensitive to outcome expectancy (Hajcak, Holroyd, Moser, & Simons, 2005; Hajcak, Moser, Holroyd, & Simons, 2007; Luu, Shane, Pratt, & Tucker, 2009; Wu & Zhou, 2009; Yeung & Sanfey, 2004). Previous research has demonstrated that the P300 is larger for unexpected relative to expected outcomes, regardless of valence. However, a few studies have found opposing evidence showing a larger P300 for unexpected gains relative to expected gains but either no difference between unexpected and expected losses (Cohen, Elger, & Ranganath, 2007) or a difference in the opposite direction (i.e., smaller P300 for unexpected loss than expected loss feedback) (Kreussel et al., 2012). Importantly, due to the temporal proximity of the P300 to the FN/RewP, there is a possibility that a large amplitude change in the P300 may have a carry-over effect on the FN/RewP that “pulls down” the amplitude of the preceding component (reviewed by (Krigolson, 2018). Due to the temporal proximity of the P200 and P300, the nature of the FN/RewP has been still clear especially in young children.

Despite many ERP studies examining FN/RewP as described above, potential sex differences in neural processing of reward-related feedback are relatively understudied in young children. However, converging evidence from both animal and human studies indicate sex differences in the development and function of the mesocorticolimbic dopamine system (Cosgrove, Mazure, & Staley, 2007), thought to support the FN/RewP (Foti, Weinberg, Dien, & Hajcak, 2011; Holroyd & Coles, 2002; Holroyd, Larsen, & Cohen, 2004). Animal studies indicate that the structure and function of the ventral tegmental area (VTA) dopaminergic system is intrinsically different in males and females (reviewed by (Gillies, Virdee, McArthur, & Dalley, 2014). The VTA – a central structure of the reward network containing mesolimbic and mesocortical dopaminergic neurons – is implicated in reward-related feedback processing and learning (Russo & Nestler, 2013). Of the few available ERP studies focusing on sex, mixed findings have been reported in older children and adults (ages 10-17 years) (Bress, Smith, Foti, Klein, & Hajcak, 2012; Crowley et al., 2013; Kujawa, Smith, Luhmann, & Hajcak, 2013).

Importantly, none of this prior work included young children (ages 4-7 years) and few studies examined the direct link between emotional negativity and ERP components relating to reward-related feedback processing (i.e., FN/RewP, P200, P300). Relatedly, existing ERP studies in adults appear to indicate sex differences in the processing of negative stimuli, with females responding more strongly to negative stimuli or unpleasant complex emotional contexts (Chentsova-Dutton & Tsai, 2007; Guillem & Mograss, 2005; Han, Fan, & Mao, 2008; Kemp, Silberstein, Armstrong, & Nathan, 2004).

The purpose of the current study was to investigate whether neural responses to reward, and their association with emotional negativity, exhibited sex-dependent effects in young children. Using ERPs, we measured neural responses to reward in children between the ages of 4-7 years. We predicted sex differences in FN/RewP amplitude at the group-level based on prior work in children (Kujawa, Proudfit, & Klein, 2014; Moser, Fisher, Hicks, Zucker, & Durbin, 2018), adolescents (Crowley et al., 2013) and adults (Yi et al., 2012). We also examined relationships between individual differences in children’s emotional negativity and ERP components during reward-related feedback processing, given prior work showing sex differences in attention allocation (Bourisly & Shuaib, 2018) and/or processing of negatively-valenced emotional stimuli (S. A. Lee, Kim, Shim, & Lee, 2017; Lithari et al., 2010). However, given mixed findings from previous studies, we did not make hypotheses about the directionality of the association between emotion negativity and the reward-related feedback ERPs.

## 2. Methods

### 2.1. Participants Recruitment and Screening

The current study included a total of one hundred forty children between the ages of four and seven. Children were recruited as part of an longitudinal study (i.e., R01MH110488) investigating relationships between neural processing of reward and early emotional development in young children with and without risk for depression. Details about recruitment and exclusion criteria were described in Supplemental Materials. For the current analyses, we focused on EEG reward processing data collected at baseline along with relevant individual difference measures. Out of n=140, n= 6 multivariate outliers detected by using the Mahalanobis Distance were excluded in major analyses. This study was approved by Duke University Institutional Review Boards and completed with the assent of child participants and the consent of parents.

### 2.2. Behavioral and Diagnostic Measures

Income-to-needs ratio: Given recent evidence suggesting that socioeconomic status is correlated with neural processing of reward (Miller, White, Chen, & Nusslock, 2021), the income-to-needs ratio was included as a covariate in regression analyses by operationalizing it as the total family income divided by the federal poverty level based on a family size (McLoyd, 1998).

2.2.2. Preschool Feelings Checklist-Scale Version (PFC-S (J. Luby, Heffelfinger, Mrakeotsky, & Hildebrand, 1999): Depression in young children has been associated with reduced neural responses to reward (Gaffrey et al., 2018). Thus, we used the PFC-S as the primary dimensional measure of depression severity for children and included it as a covariate in regression analyses. It is a validated measure of depression severity in 3-7-year-olds (J. Luby, Lenze, & Tillman, 2012; J. L. Luby, Heffelfinger, Koenig-McNaught, Brown, & Spitznagel, 2004). It consists of 23 items adapted from the Preschool Feelings Checklist (J. L. Luby et al., 2004) (PFC).

2.2.3 Beck Depression Inventory-2 (BDI; (Beck, Steer, Ball, & Ranieri, 1996): The BDI is a well validated 21-item self-report inventory for adults measuring depression symptomatology. Recent studies indicate that parental depression may lead to inflate parent-reported child’s negative emotionality and/or risk for psychopathology (Bouvette-Turcot et al., 2020; Merwin, Smith, Kushner, Lemay, & Dougherty, 2017). Thus, to control for potential inflation of parent-reported child’s depression or negative emotionality, we used BDI-2 as another covariate in our major analyses.

2.2.4. The Emotion Regulation Checklist (ERC; (Shields & Cicchetti, 1997): It is a parent report measure of child emotion-regulation (Shields, Dickstein et al. 2001). It includes both positively and negatively weighted items rated on a 4-point Likert scale that can be used to produce Lability/Negativity and Emotion Regulation subscale scores. Emotion Regulation scores reflect the expression of emotions, empathy and emotional self-awareness, while the Negativity scores reflect a lack of flexibility, anger dysregulation, and mood lability. In current study, we focused on the Negativity scores of the ERC to measure individual differences in child’s emotional negativity and its relation to reward-related feedback processing.

### 2.3. EEG Task and Materials

A developmentally sensitive version of the Doors Guessing Task based on previous research was used to measure neural responses to reward (Foti et al., 2011; Gaffrey et al., 2018). Briefly, at the beginning of each trial, participants were presented with an image of two doors. Children were told that there is candy hiding behind one of the doors. Given the young age of the children participating in the study, a one button choice strategy was used. That is, one door had a button on it while the other did not. Children were asked to choose the door with the button on it by pressing a button or to choose the other door without the button by not doing anything (i.e., not pressing). Which door had the button on it, as well as its association with a win or loss outcome, was pseudorandomized to prevent any specific pattern of wins/losses based on button pressing. Candy prizes (i.e., win: get four skittles, loss: lose two skittles) instead of the typical monetary incentives were used since young children do not have a clear conception of money (Hennefield et al., 2021) and react positively to prizes and gifts (Luking, Luby, & Barch, 2014). The task was administered through E-Prime presentation software (Psychology Software Tools, Inc, Sharpsburg, PA). Following previous research (Belden et al., 2016; Tunison, Sylvain, Sterr, Hiley, & Carlson, 2019), we focused on comparing reward gain and loss outcomes only in the current study analyses (see Figure 1 for the task paradigm).

**Figure 1.**
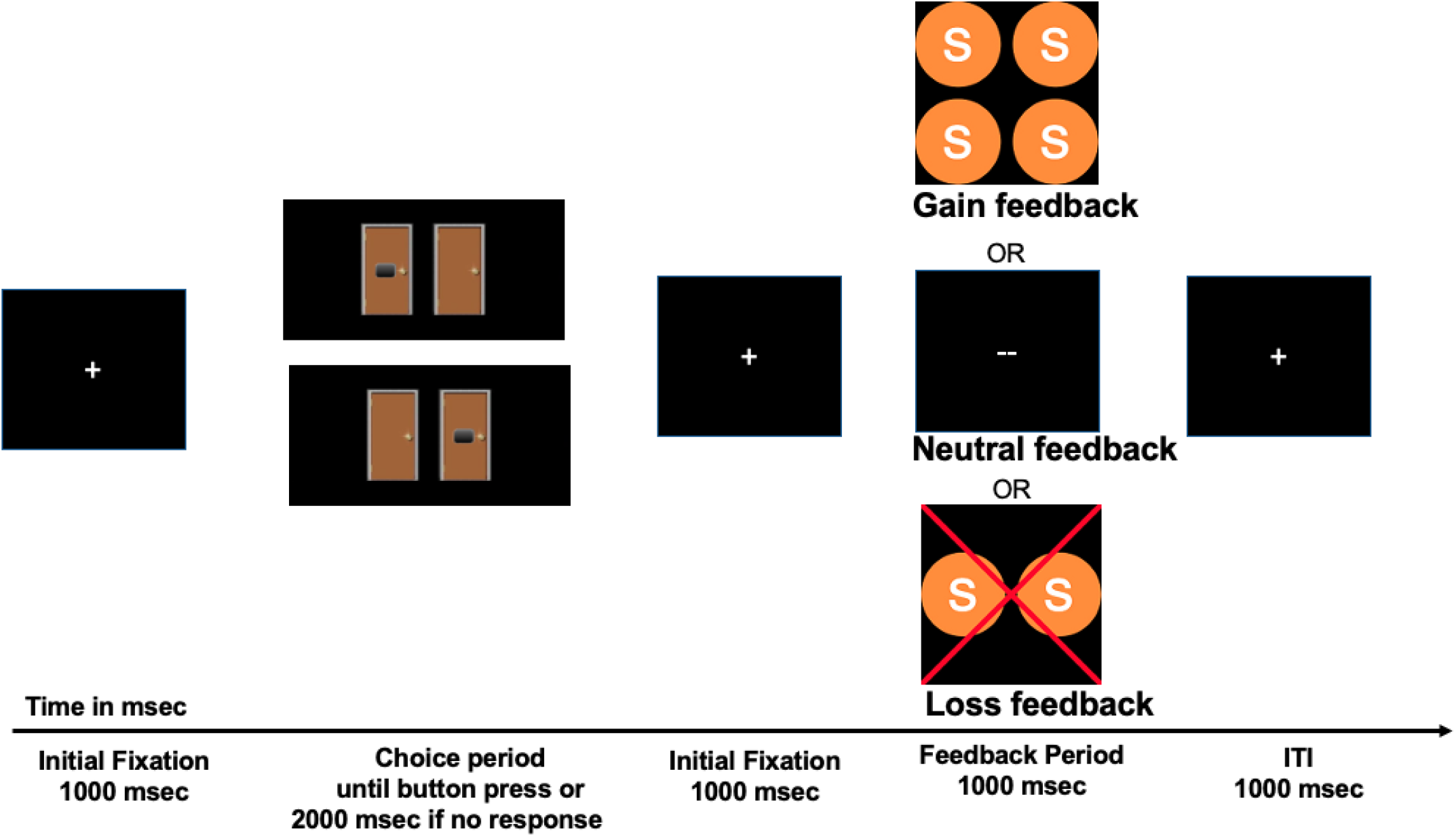
The Modified Doors Task (^1,2^) *Note*. If children chose the ‘correct’ door, they would see four pieces of candy. If they chose the wrong door, they would see two candies with a red ‘X’ through it. There was also the possibility of a neutral outcome occurring where children neither won or lost candy, illustrated as two dashes appearing on the outcome screen. The image of the doors remained on the screen until the participants chose the door with the button or for 2 seconds if no button press occurred. We focused on gain and loss feedback only due to different physiological features of neutral feedback in current study.

### 2.4. EEG data acquisition and preprocessing

Continuous EEG was recorded using EGI’s Net Station acquisition software version 5.4.2 with the128-channel Geodesic Sensor Net system at a 1000 Hertz (Hz) sampling frequency. Data was referenced online to the vertex (Cz) and impedances were maintained below 50 kilohms throughout the paradigm.

Offline EEG data preprocessing was performed using EEGLAB version 2021 (Delorme & Makeig, 2004) in MATLAB R2019b (The MathWorks Inc, Natick, MA) using custom scripts available at https://github.com/DEEDLabEEG. First, twenty-four channels located on the outer ring of the sensor net were removed because they contained many artifacts due to their location close to the neck and on the face. The data were then downsampled to 250Hz and segments of data without relevant events (i.e., breaks) were removed. Using the ERPLAB v8.2 plugin (Lopez-Calderon & Luck, 2014), a 0.1 to 30 Hz, 4^th^ order Butterworth, bandpass filter was applied. CleanLine (T. Mullen, 2012) was used to remove any remaining 60 Hz line noise (Mitra & Bokil, 2007). All data were re-referenced to the average of the mastoids. Bad channels were removed by running the Clean Rawdata plugin; a channel was considered bad if 1) it was flat for more than five seconds, 2) contained more than four standard deviations of line noise relative to its signal, or 3) correlated at less than .8 to nearby channels.

A copy of the data was created and high-pass filtered at 1 Hz (4^th^ order Butterworth) using EEGLAB v2021 (Delorme & Makeig, 2004). Artifact Subspace Reconstruction (ASR; (T. R. Mullen et al., 2015) with the burst criterion set to 20 (Chang, Hsu, Pion-Tonachini, & Jung, 2018) was run to remove large artifacts with Clean Rawdata. Extended infomax independent component analysis (ICA; (T. W. Lee, Girolami, & Sejnowski, 1999) with PCA dimension reduction (50 components) was run and the resulting ICA matrix was then copied over to the original full-length data (i.e., the data just before the copy was made and the 1 Hz high-pass filter was applied). The ICLabel plugin (Pion-Tonachini, Kreutz-Delgado, & Makeig, 2019) automatically removed independent components with a probability greater than .7 of being associated with eye movements and blinks. Application of the ICA matrix to the full-length data and subsequent removal of eye-related components allowed for the preservation of data that would have otherwise been removed by ASR or other artifact removal methods. Bad channels that were previously removed were interpolated using spherical splines.

The EEG was segmented for each gain and loss trial, beginning 500 ms before feedback onset and continuous for 800ms following feedback onset. Baseline correction was applied to the 200ms window before feedback onset. Artifact rejection using the Trial-By-Trial plugin (Ben-Shachar, 2018) removed epochs with at least 10 channels meeting the following criteria: 1) peak-to-peak amplitudes exceeding 100 μV within 200 ms windows sliding by 20 ms, 2) voltages less than -150 or greater than +150 μV, or 3) joint probabilities above three standard deviations for local and global thresholds. If less than 10 channels in an epoch met criteria for rejection, the epoch was not removed, but the identified channels were interpolated for that epoch only. Lastly, the averaged ERPs for each bin (i.e., gain, loss) were computed. The mean of the number of accepted gain and loss trials were 24.57(±5.25) and 24.02 (±5.56), respectively.

### 2.5. Temporospatial Principal Components Analysis (PCA)

Previous research suggests that the RewP/FN may overlap with other reward-related processing ERP components, including the P200 and P300 (Schaefer, Buratto, Goto, & Brotherhood, 2016; Schuermann, Endrass, & Kathmann, 2012). Relatedly, research has suggested that using a data reduction technique such as PCA can successfully decompose a raw ERP waveform across time into its underlying components (Donchin E & E, 1979), allowing for the isolation of the RewP/FN from other temporally or spatially overlapping components.

Following procedures used in prior work (Foti et al., 2011), we conducted the temporospatial PCA as follows on the averaged baseline-correct ERPs for gain and loss trials separately. The RewP/FN was quantified using temporospatial PCA, a technique that extracts linear combinations of data points using criteria distinguishing between consistent patterns of electrocortical activity (Dien & Frishkoff, 2005). This analysis was conducted using the ERP PCA Toolkit, version 2.93 (Dien, 2010a). Following prior guidelines for applying PCA to ERP datasets (Dien, 2010b; Dien, Khoe, & Mangun, 2007), a temporal PCA was performed on the data first to capture variance across time points. Promax rotation was used, and 18 temporal factors were extracted based on the resulting Scree plot (Cattell, 1966). For each temporal factor, this analysis yielded factor scores for each combination of electrode, participant, and trial type, representing the amount of activity in the original data captured by that factor. The spatial distribution of these factor scores was then analyzed using spatial PCA. A separate spatial PCA was performed for each of the 18 temporal factors with Infomax rotation. Based on the averaged Scree plot for all 18 temporal factors, 5 spatial factors were extracted, yielding 90 unique factors combinations. The covariance matrix and Kaiser normalization were used for each PCA. The waveforms for each factor were reconstructed (i.e., converted to microvolts) by multiplying the factor pattern matrix with the standard deviations.

### 2.6. Statistical analyses

Statistical analyses were conducted using IBM SPSS Statistics (IBM Corp, Armonk, NY; Version 27). The FN and RewP were defined as the mean amplitude between 344 and 444 ms following feedback indicating loss or gain, respectively, from each PCA factor representing the ERP of interest. The identified mean amplitudes from loss or gain for each individual were subsequently exported to SPSS for use in linear regressions. Residualized scores of loss using regression (i.e., FN: unstandardized residuals of loss controlling for gain) or gain (i.e., RewP: unstandardized residuals of RewP controlling for loss) were calculated in SPSS. Similarly, P200 and P300 were calculated as residualized gain scores, but the mean amplitude of P200 was measured between 192 and 292 ms while the same time window of 344-444 ms following feedback indicating gain was used for P300 like the RewP/FN.

Sex differences in age, parental depression, and child depression and emotional negativity scores were examined using independent *t*-test (see Table 1). Hierarchical regressions were used to investigate whether main effect of sex and/or sex by emotional negativity interaction would predict RewP, P200, and P300 relating to gain or relating to loss while controlling for income-to-need ratio, parental or child depression.

**Table 1.**
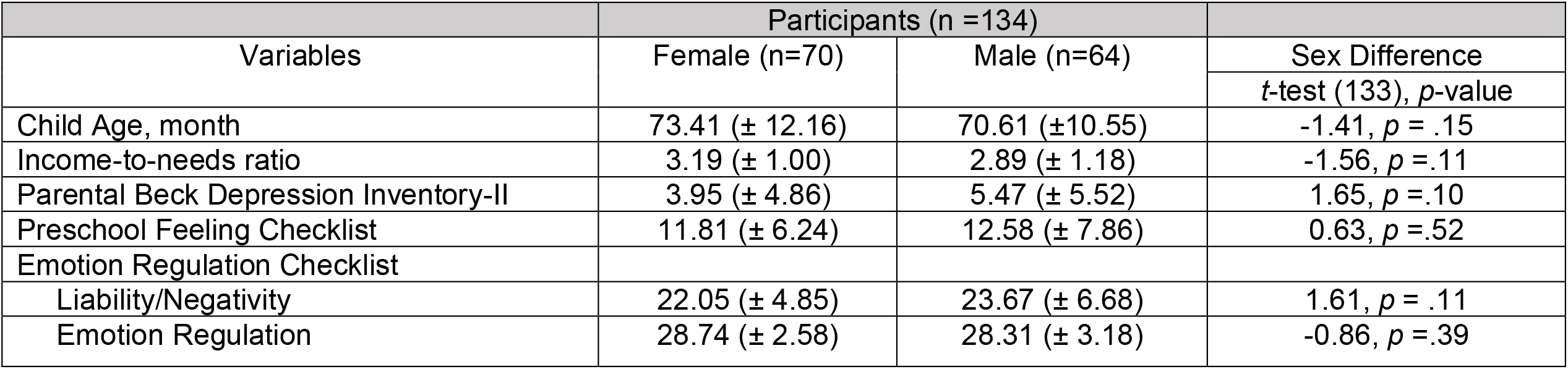
Participants Characteristics

| Variables | Participants (n =134) |  | Sex Difference |
| --- | --- | --- | --- |
|  | Female (n=70) | Male (n=64) | *t*-test (133), *p*-value |
| Child Age, month | 73.41 (± 12.16) | 70.61 (±10.55) | -1.41, *p* = .15 |
| Income-to-needs ratio | 3.19 (± 1.00) | 2.89 (± 1.18) | -1.56, *p* =.11 |
| Parental Beck Depression Inventory-II | 3.95 (± 4.86) | 5.47 (± 5.52) | 1.65, *p* =.10 |
| Preschool Feeling Checklist | 11.81 (± 6.24) | 12.58 (± 7.86) | 0.63, *p* =.52 |
| Emotion Regulation Checklist |  |  |  |
| Liability/Negativity | 22.05 (± 4.85) | 23.67 (± 6.68) | 1.61, *p* = .11 |
| Emotion Regulation | 28.74 (± 2.58) | 28.31 (± 3.18) | -0.86, *p* =.39 |

## 3. Results

As presented in Table 1, males and females did not show any sex differences in age and other individual difference measures of depression and emotionality.

### 3.1. The grand average waveforms before PCA

Waveforms depicting the grand average ERPs (i.e., before PCA) are presented in Figure 2. Consistent with prior work with adults (Foti et al., 2011; Proudfit, 2015), our data showed the presence of FN (i.e., loss-gain) or RewP (i.e., gain-loss) at frontocentral electrodes including E19 in young children. In addition, and again consistent with prior work in older age groups, we confirmed that the deflection in the waveform for gain minus loss was superimposed on the P300 at ∼400ms.

**Figure 2.**
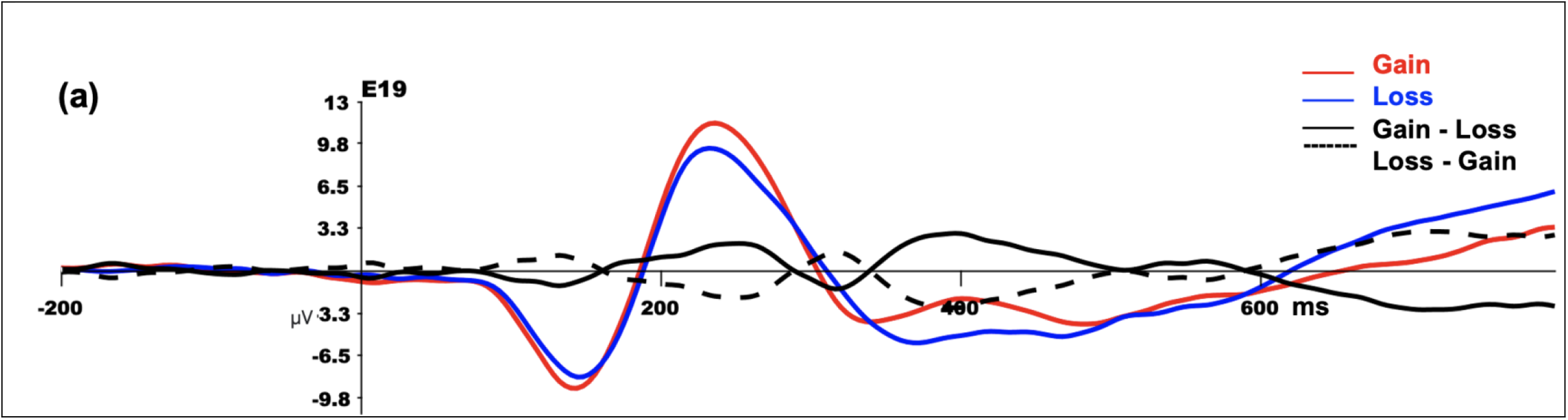
a) The original grand average waveform of the reward positivity at EEG19 for loss and gain difference waveforms across all subjects before the PCA.

When we explored potential sex differences in gain-related RewP or loss-related FN amplitudes *before* the PCA method, no sex differences were observed [*t*(131) = 1.30, *p* = .19; *t*(131) = -1.30, *p*=.20, respectively)].

#### PCA results

Of the 90 total factor combinations yielded by the PCA, 18 accounted for at least 1% of the total variance in the data. Consistent with prior work by (Dien & Frishkoff, 2005), visual inspection of the waveforms associated with these 18 factors was used to select those that most readily corresponded to ERP components relevant to the current task paradigm. Four factors were chosen for further statistical analysis, corresponding to the P200, RewP, FN, and P300 (Table 2 and Figure 3 for each PCA factor representing each ERP of interest and related waveforms and topography, respectively).

**Table 2.** PCA factor combinations selected for statistical analysis

| Corresponding ERP component | Temporospatial factor combination | Variance explained (%) | Temporal loading peak (ms) | Spatial distribution | Peak channel | Gain (mean (SD)) | Loss (mean (SD)) | Gain - Loss $t(133)$ |
| --- | --- | --- | --- | --- | --- | --- | --- | --- |
| P200 | TF4/SF1 | 3.7 | 240 | Fronto-central positivity | EEG 6 | 6.59 (5.06) | 4.87 (5.09) | 5.03*** |
| RewP or FN<sup>a</sup> | TF1/SF1 | 15 | 392 | Fronto-central negativity | EEG19 | -2.20 (8.33) | -3.53 (7.81) | 2.17* |
| P300 | TF1/SF2 | 5.4 | 392 | Centroparietal positivity | EEG 71 | 6.63 (5.99) | 7.44 (6.76) | -1.56 |

Note. FN = Feedback-related Negativity, RewP = Reward-related positivity,
<sup>a</sup> The FN amplitudes, calculated by loss - gain contrast were significant different ( $p = .03$ )
\* $p = .03$ , \*\*\* $p < .001$

**Figure 3.**
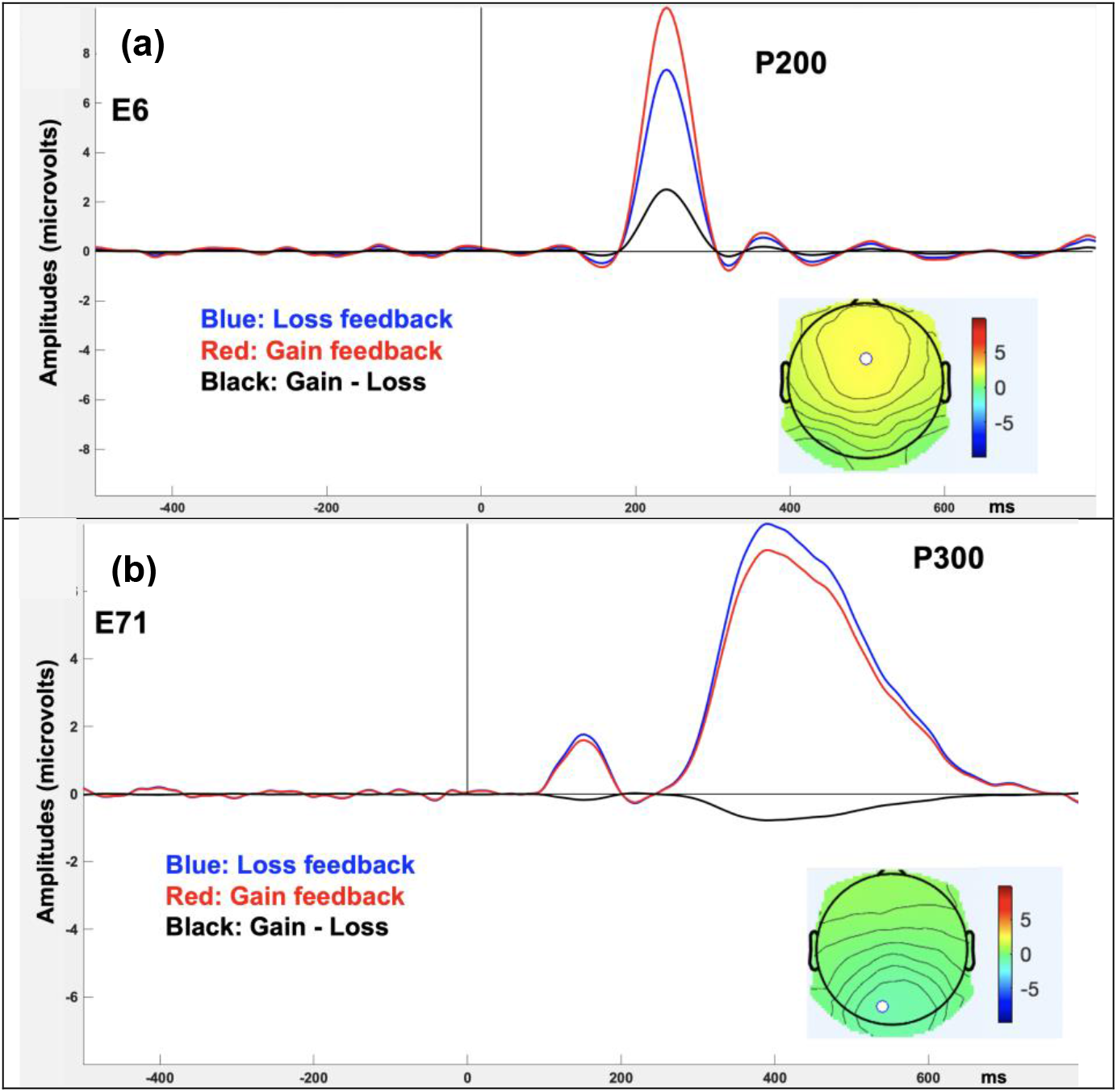
P200 and P300 after PCA analysis. Notes. (a) The waveforms at EEG 6 representing the portion of the ERP associated with TF4/SF1, the PCA factor corresponding to the P200 ERP component, (b) The waveforms at EEG 71 representing the portion of the ERP associated with TF1/SF2, the PCA factor corresponding to the P300 ERP component. Topographies presented in (a), and (b) represents the scalp topography of the difference between gain and loss trials with white dot indicating the location of each EEG electrode when temporal loading is maximal

The waveforms and scalp topography associated with Temporal Factor 4, Spatial Factor 1 (TF4/SF1), the PCA factor corresponding to the P200 is presented in Figure 3.a. As evident in the grand average ERP waveforms of the PCA, this PCA factor represents a relative positivity for gain compared to loss trials that is maximal at frontocentral sites with a peak site of EEG 6, with the difference across feedback types being statistically significant (Table 2). The waveforms and scalp topography associated with Temporal Factor 1, Spatial Factor 2 (TF1/SF2), the PCA factor corresponding to the P300 (i.e., gain-loss) is presented in Figure 3.b. As presented in the grand average ERP waveforms of the PCA, this PCA factor shows similar degree of ERP amplitudes during gain versus loss trials that is maximal at centroparietal sites with a peak site of EEG 71, with the no statistical differences across feedback types (Table 2). Lastly, the waveforms and scalp topography associated with Temporal Factor 1, Spatial Factor 1 (TF1/SF1), the PCA factor corresponding to the RewP (i.e., gain-loss) is presented in Figure 4. As evident in the grand average ERP waveforms of the PCA, this PCA factor represents a relative positivity for gain compared to loss trials that is maximal at frontocentral sites with a peak site of EEG 19, with the difference across feedback types being statistically significant (Table 2). Interestingly, the PCA waveform reveals negative responses to both gain and loss trials, with the less negative response for gain resulting in a positivity when compared to loss.

**Figure 4.**
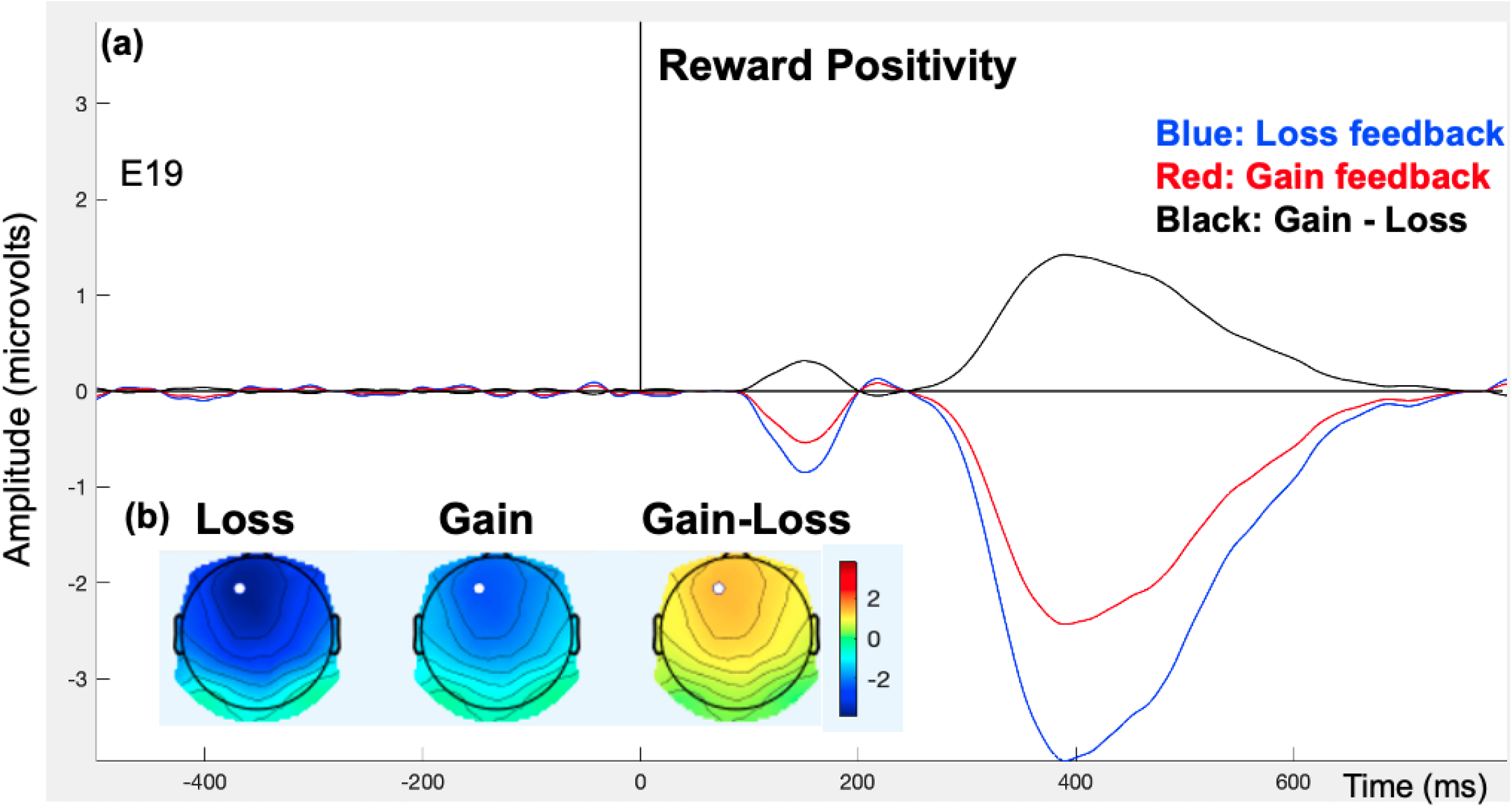
a) The waveforms at EEG 19 representing the portion of the ERP associated with TF1/SF1, the PCA factor corresponding to the reward positivity. B) The scalp topography of the difference between gain and loss trials at 344-444 ms, where the temporal loading is maximal. White dot indicates the location of EEG 19, peak site.

### 3.2. Sex differences in PCA-derived FN, P200, and P300 amplitudes

We conducted independent *t*-tests to examine sex differences in PCA-driven ERP components showing significant differences in neural responses during gain versus loss (i.e., P200, RewP) using each temporal and spatial factor combinations generated from the PCA. Males showed greater loss-related FN amplitudes from TF1/SF1 factor combination [(*t*)131) = -2.43, *p*=.02, Cohen’s *d* =0.25] but there were no sex differences in gain-related RewP amplitudes from T1/SF1 factor combination, (*t*)128) = 1.47, *p*=.14]. Regarding PCA-driven P200, males showed both greater gain-related P200 from TF4SF1 [(*t*)131) = 3.32, *p*=.001, Cohen’s *d* = 0.57] and greater loss-related P200 from TF4SF1[(*t*)131) = -2.17, *p*=.03, Cohen’s *d* =0.37] compared to females.

### 3.3. Hierarchical regression

As presented in Table 3, while controlling for age, income-to-needs ratio, parental and child’s depression severity scores, hierarchical regression revealed that the interaction of sex and negativity, as measured by subscale of Liability/Negativity from the ERC, predicted gain-related P200 (i.e., residualized gain in P200). According to *post hoc* correlations for each sex, greater negative emotionality scores for males only were associated with the less P200 amplitudes during gain (*r* = -0.32, *p* =.01, *n* = 64) while females did not show such association (*r* = -0.07, *p* =.54, *n* = 67) (Figure 5).

**Table 3.** Summary of linear regression analysis for variables predicting P200 relating to Gain

| | | *B* | *SE B* | $\beta$ | *t* | Significance | *R*<sup>2</sup> change | Sig. *F* change |
| --- | --- | --- | --- | --- | --- | --- | --- | --- |
| Step 1 |  |  |  |  |  |  | 0.08 | .04 |
|  | Child age | -.04 | .02 | -.13 | -1.47 | .14 |  |  |
|  | Income-to-needs ratio | .07 | .29 | .02 | 0.24 | .80 |  |  |
|  | Sex | 1.72 | .63 | .24 | 2.73 | .007 |  |  |
|  | Parental depression | -.01 | .06 | -.02 | -0.27 | .78 |  |  |
|  | Child depression | -.03 | .04 | -.07 | -0.83 | .40 |  |  |
| Step 2 |  |  |  |  |  |  | .03 | .06 |
|  | Negativity | -.12 | .06 | -.20 | -1.88 | .06 |  |  |
| Step 3 |  |  |  |  |  |  | .04 | .02 |
|  | Sex by Negativity | -.25 | .10 | -.86 | -2.37 | .01 |  |  |

**Figure 5.**
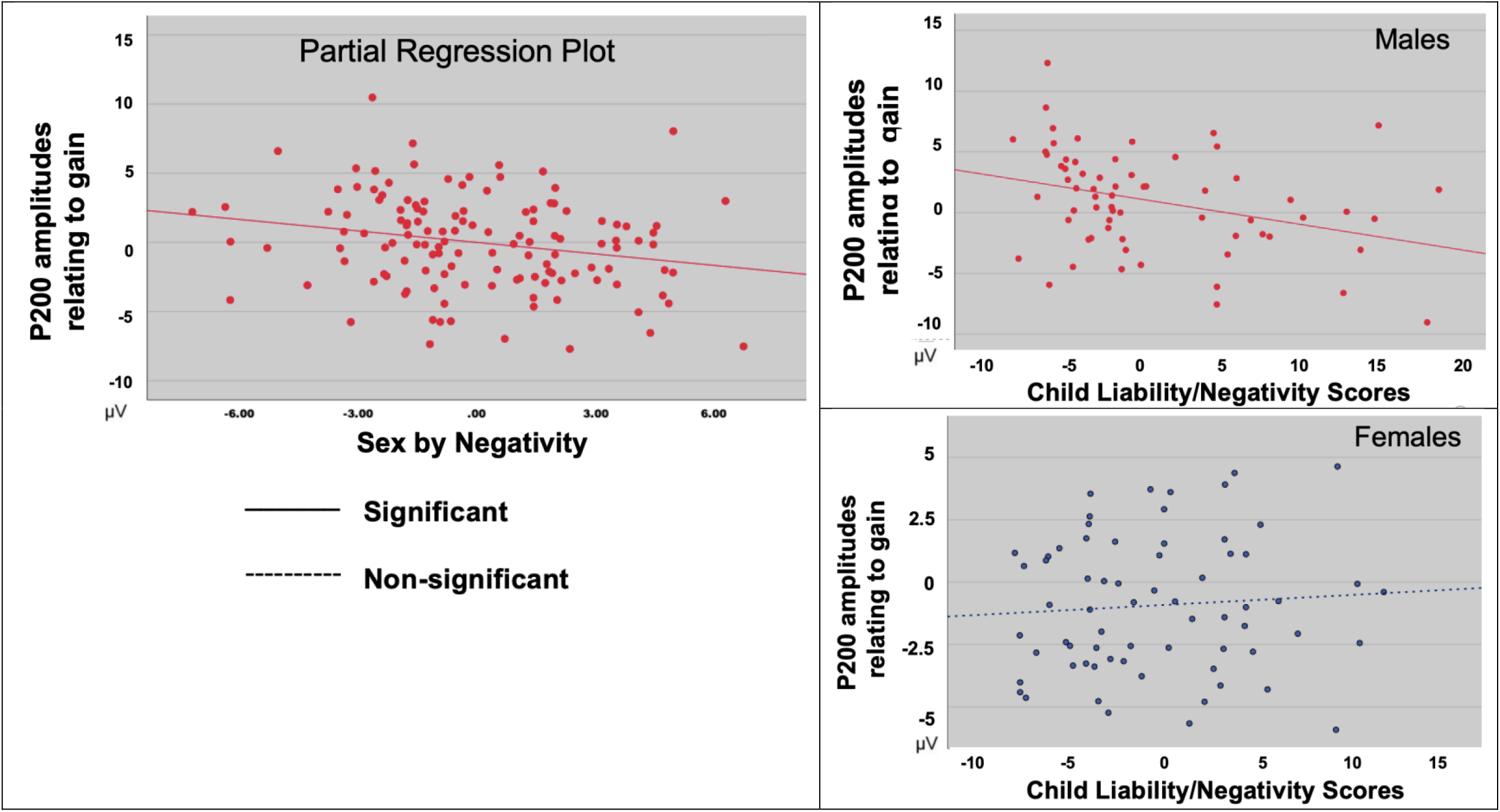
Males only show negative association between child negativity and residualized P200 relating to gain feedback.

We also conducted the same regression models with loss-related FN and loss-related P200 amplitudes as dependent variables but did not find any significant models (all *P*s > .16).

## 4. Discussion

The current study investigated the associations between neural responses to reward, emotion, and biological sex in 4-7-year-olds using ERPs. Importantly, using temporospatial PCA, we were able to successfully separate and identify the RewP/FN, P200, and P300 in our sample of young children. As a result, at the group level, we were able to investigate each ERP separately and found greater loss-related FN amplitudes, and greater gain- and loss-related P200 amplitudes in males relative to females. In addition, we also found that emotional negativity correlated with the P200 in males only, suggesting that male children with higher emotional negativity may pay less attention to gain feedback or find it less salient. As a result, the current findings suggest that neural processing of reward and its association with emotional negativity may already differ based on biological sex during early childhood.

Sex differences in loss-related FN and reward-related P200 ERP components in young children As hypothesized, we found sex differences in neural processing of reward feedback even in young children with small-to-medium effect sizes. Males showed greater FN amplitudes relating to loss feedback compared to females at a group level, while both showed similar levels of RewP amplitudes relating to gain feedback. Existing ERP studies that examined sex differences in reward processing have reported mixed findings across developmental stages and task paradigms (e.g., (Bress et al., 2012). One possible factor contributing to mixed findings is that the RewP (or FN) ERP component overlaps in time with other ERP components, potentially obscuring sex differences. Supporting this possibility, we did not see sex differences in gain-related RewP or loss-related FN *before* the PCA method was applied. Nonetheless, the current findings of sex differences in young children during loss feedback processing are in line with some prior ERP studies. For example, (Kujawa et al., 2013) found that males had a larger FN amplitudes than females (ages 18-27years). (Ding et al., 2017) using children (ages 9-12 years) also found sex differences in electrophysiological responses both to reward gain and loss feedback; males showed less FN amplitudes under gain and loss conditions, and longer FN latency in gain and loss conditions than females. Further, (Crowley et al., 2013) investigated developmental changes in the patterns of the FN amplitudes and latencies with a relatively small number of participants in each developmental stage (*n* = 91, ages 10-17 years). While our study found sex differences only in the loss-related FN amplitudes, (Crowley et al., 2013)’s work revealed that males showed larger responses irrespective of feedback type and longer latencies for rewarded feedback during late childhood to middle adolescence. Future studies with large sample sizes and using the PCA method are required to confirm whether sex differences in neural processing of reward feedback are specific to childhood only or more general to different developmental stages.

Similar to findings of sex differences in the FN ERP findings, we also found that males exhibited greater gain-related and loss-related P200 amplitudes relative to females at a group level. Despite emerging evidence suggesting sex differences in attention during processing of reward gain or loss (Dumais, Chernyak, Nickerson, & Janes, 2018; Greimel et al., 2018; Pfabigan et al., 2014) or negative stimuli (Jin et al., 2013; S. A. Lee et al., 2017), the majority of ERP studies in adults did not include sex as a variable in their analyses (e.g., reviewed by (Torrence & Troup, 2018). However, our current findings of sex differences in the P200 in young children are in line with some prior work in adults, suggesting that the P200 is affected by sex; although the directionality of sex differences in P200 amplitudes is inconsistent across those studies (Bourisly & Shuaib, 2018; Lijffijt et al., 2009; Xin, Gu, Wang, Lei, & Li, 2021; Yuan, He, Qinglin, Chen, & Li, 2008). For example, (Yuan et al., 2008) found that men (*n* = 15, mean age = 20.8 years) exhibited smaller amplitudes and longer latencies than women (*n* = 15, mean age = 21.1 years) for the P200 during an oddball task, a commonly used task for cognitive and attention measurement in ERP studies. Contrary to this, (Bourisly & Shuaib, 2018) found that males (*n* = 15, mean age = 39 years) had a significantly larger P200 peak amplitude and a significantly shorter P200 latency period relative to females (*n* = 15, mean age = 40.6 years) during a similar type of oddball task. Many studies indicate that efficient attention allocation towards rewarded stimuli or reward-predicting stimuli is a critical component of reward-based learning (e.g., (Tibboel & Liefooghe, 2020; Vartak, Jeurissen, Self, & Roelfsema, 2017). In addition, reduced P200 amplitudes have been associated with learning difficulties in reading, writing, and arithmetic domains in children (Fernández, Silva-Pereyra, Prieto-Corona, Rodríguez-Camacho, & Reynoso-Alcántara, 2014). Considering that dysfunctional reward-based learning has been implicated in many neurodevelopmental disorders showing sex differences (e.g., substance use: (Dichter, Damiano, & Allen, 2012), further investigating whether the P200 supports reward-based learning behaviors in a sex-specific way is an important next step.

Sex-specific association between gain-related P200 ERP and individual differences in emotional negativity in young children Understanding the nature of feedback processing in young children is not only critical to developing appropriate educational guidelines for children’s self-regulated learning (reviewed in (Panadero, 2017), but also important to better understand the developmental trajectories of socioemotional development. In addition to group-level sex differences, our study also identified sex-related differences in association between P200 amplitude to gain feedback and child’s emotional negativity. That is, our results showed that only males having higher emotional negativity tended to show smaller P200 amplitudes relating to gain feedback, potentially reflecting less attention towards rewarding stimuli. Failure of attention shaping as a function of reward has been considered as one key characteristics of depression (Keller, Leikauf, Holt-Gosselin, Staveland, & Williams, 2019) and depression risk (Vrijen, Hartman, & Oldehinkel, 2019). For example, difficulties in shifting attention towards expected rewards at 16 years predicted depression during 3-year later follow-up assessment (Vrijen et al., 2019). Current results suggest that this early relationship between less attention towards reward gain (i.e., reduced P200) and higher emotional negativity may reflect a potential biomarker of altered socioemotional development in young males. Future longitudinal studies are needed to investigate how this early relationship may predict later onset of psychopathology such as depression in a sex-specific way.

Our study also revealed sex-specific relevance of the P200 amplitudes relating to gain feedback and child’s emotional negativity. That is, our results showed that only males having higher emotional negativity tended to show smaller P200 amplitudes relating to gain feedback, reflecting less attention towards rewarding stimuli. Mounting evidence suggests that emotional negativity such as irritability is a sensitive marker of risk for later depression in children (as early as three years old)(Brotman, Kircanski, Stringaris, Pine, & Leibenluft, 2017; J. L. Luby et al., 2003). One example of emotional negativity, irritability characterizes it as excessive reactivity to negative emotional stimuli (Caprara GV et al., 1985; L., 1993). Several longitudinal studies show that irritability-related traits in nonclinical populations during childhood are predictors of psychopathology including depression. The existing but limited empirical data suggests that negatively valenced social and emotional stimuli may be particularly salient for irritable youths and thus more likely to capture their attention (reviewed in (Leibenluft & Stoddard, 2013). In line with these, current results suggest that this early relationship between less attention towards reward gain (i.e., reduced P200) and higher emotional negativity may reflect a potential biomarker of altered socioemotional development in young males. Future longitudinal studies are needed to investigate how this early relationship may predict later onset of psychopathology such as depression in a sex-specific way.

We did not find sex-specific associations between RewP or FN and emotional negativity in young children, which was surprising. Because we expected to find at least the link of RewP or FN amplitudes and emotional negativity. Several ERP studies in adults appear to indicate that gain-related RewP ERP is modulated by hedonic “liking”, and both state and trait affect(Angus, Kemkes, Schutter, & Harmon-Jones, 2015; Brown, Jackson, & Cavanagh, 2022). However, we do not know such relationship was sex-specific or not, as most of those study did not include sex as an independent variable. We speculate that potential sex-specific developmental trajectories of reward processing might be driven by different attention allocation between the sexes, as indexed by early P200 amplitudes rather than later feedback processing itself. To support this, several studies have found sex differences in the developmental neural networks supporting attention (Dumais et al., 2018; Gaillard, Fehring, & Rossell, 2021; Rubia, Hyde, Halari, Giampietro, & Smith, 2010) but this should be confirmed in future studies.

### Limitations

Although our study provided novel findings of sex-related effects for both group-level and individual-level factors related to neural processing of feedback in young children, there are still unanswered questions that should be tested in future studies. First, while the current task paradigm (i.e., the modified doors task) has been used commonly as reward processing task in children (e.g. (Belden et al., 2016; Hennefield et al., 2021), it did not include a reward anticipation phase but only reward receipt and/or feedback processing. Emerging evidence clearly indicates that anhedonia, emerging as a highly promising endophenotype of depression, may be associated with less reward anticipation rather than reward-related feedback processing itself (Rizvi, Pizzagalli, Sproule, & Kennedy, 2016; Sherdell, Waugh, & Gotlib, 2012). In addition, it is not clear whether sex-related effects found in the current study might be specific to young childhood or generalized to later development of feedback processing in older children and adolescents. Longitudinal study designs would be ideal to replicate sex-related effects on reward-related feedback processing across developmental stages. Second, although several studies localized the anterior cingulate cortex (ACC) and/or striatum as potential neural generators of the RewP/FN in adults (Foti et al., 2011; Proudfit, 2015), it is necessary to confirm whether or not the same mesocorticolimbic functional connectivity with the ACC are implicated in supporting the RewP/FN ERPs in young children. Lastly, our sample has been limited due to the lack of diversity. Larger sample with various socioeconomic status might have revealed different patterns of sex differences and/or sex-specific association between P200 or RewP and emotional negativity (Perera, Salehuddin, Khairudin, & Schaefer, 2021).

## Conclusions

Using EEG to investigate neural reward processing in young children, we report novel sex-specific effects at group and individual levels. Specifically, we identified greater loss-related FN and gain- and loss-related P200 amplitudes in males relative to females and an association between gain-related P200 amplitudes and emotional negativity for males only. These findings suggest that sex-specific differences in the early development of neural reward processing are already present in young children. Future longitudinal research in young children investigating sex-related differences in the neural correlates of reward processing are highly likely to provide valuable information for preventing or mitigating potential sex differences in altered reward-related behavior (e.g., drug use) and/or psychopathology (e.g., depression) at later ages.

