## Supplemental Materials for "Sex-specific association between emotion negativity and neural processing of reward feedback in young children"

Methods

Children and their families were recruited from the Research Participation at Duke Database maintained by the Human Subjects Coordinator for the Department of Psychology and Neuroscience at Duke University and/or from local community events. Parents initially completed the Preschool Feelings Checklist (PFC), a validated screening measure for elevated symptoms and risk for depression in 3-7-year-olds (Luby, Heffelfinger, Mrakeotsky, & Hildebrand, 1999). Following our previous work (Gaffrey et al., 2018), parents endorsing greater than 3 items (high risk) or 1 item or less (low risk) were invited for further eligibility screening via a telephone call.

Children without exclusions were invited to participate in the full study. Exclusion criteria were as follows; IQ < 70, developmental delay, premature birth (< 36weeks gestation), neurological condition, substance exposure in utero, or psychiatric medication. Parents and children were compensated for their time.
